# Noncoding microdeletion in mouse *Hgf* disrupts neural crest migration into the stria vascularis, reduces the endocochlear potential and suggests the neuropathology for human nonsyndromic deafness DFNB39

**DOI:** 10.1101/778365

**Authors:** Robert J. Morell, Rafal Olszewski, Risa Tona, Samuel Leitess, Julie M. Schultz, Elizabeth J. Thomason, Brittany N. Whitley, Connor Hill, Thomas Saunders, Matthew F. Starost, Tracy Fitzgerald, Elizabeth Wilson, Takahiro Ohyama, Thomas B. Friedman, Michael Hoa

## Abstract

Hepatocyte growth factor (HGF) is a multifunctional protein that signals through the MET receptor. HGF stimulates cell proliferation, cell dispersion, neuronal survival and wound healing. In the inner ear, levels of HGF must be fine-tuned for normal hearing. In mouse, a deficiency of HGF expression limited to the auditory system, or over-expression of HGF, cause neurosensory deafness. In human, noncoding variants in *HGF* are associated with nonsyndromic deafness *DFNB39*. However, the mechanism by which these noncoding variants causes deafness was unknown. Here, we reveal the cause of this deafness using a mouse model engineered with a noncoding intronic 10bp deletion (del10) in *Hgf*, which is located in the 3’UTR of a conserved short isoform (*Hgf/NK0.5*). Mice homozygous for del10 exhibit moderate-to-profound hearing loss at four weeks of age as measured by pure-tone auditory brainstem responses (ABRs). The wild type +80 millivolt endocochlear potential (EP) was significantly reduced in homozygous del10 mice compared to wild type littermates. In normal cochlea, EPs are dependent on ion homeostasis mediated by the stria vascularis (SV). Previous studies showed that developmental incorporation of neural crest cells into the SV depends on signaling from HGF/MET. We show by immunohistochemistry that in del10 homozygotes, neural crest cells fail to infiltrate the developing SV intermediate layer. Phenotyping and RNAseq analyses reveal no other significant abnormalities in other tissues. We conclude that, in the inner ear, the noncoding del10 mutation in *Hgf* leads to dysfunctional ion homeostasis in the SV and a loss of EP, recapitulating human DFNB39 deafness.

**Significance Statement:** Hereditary deafness is a common, clinically and genetically heterogeneous neurosensory disorder. Previously we reported that human deafness DFNB39 is associated with noncoding variants in the 3’UTR of a short isoform of *HGF* encoding hepatocyte growth factor. For normal hearing, HGF levels must be fined-tuned as an excess or deficiency of HGF cause deafness in mouse. Using a *Hgf* mutant mouse with a small 10 base pair deletion recapitulating a human DFNB39 noncoding variant, we demonstrate that neural crest cells fail to migrate into the stria vascularis intermediate layer, resulting in a significantly reduced endocochlear potential, the driving force for sound transduction by inner ear hair cells. HGF-associated deafness is a neurocristopathy but, unlike many other neurocristopathies, it is not syndromic.

## Introduction

Hepatocyte growth factor (HGF) is an activator of mitosis and identical to “scatter factor”, which stimulates epithelial cells to disperse in culture (Stoker et al., 1987; Nakamura, 1989). HGF is also implicated in branching morphogenesis (Zhang and Vande Woude, 2003), tumorigenesis (Zhang et al., 2018), immune cell regulation (Papaccio et al., 2018), neuronal survival (Thompson et al., 2004; Nakano et al., 2017), wound healing (Miyagi et al., 2018), neuronal differentiation, synapse formation and maturation (Matsumoto et al., 2014). There are numerous studies of HGF structure, domains, splice isoforms and diverse functions (Comoglio et al., 2003; Matsumoto et al., 2014). Inactive pre-pro-HGF is secreted and proteolytically processed into a functional α and β disulfide-linked heterodimer (Fig. 1*A*). The α-chain contains an N-terminal hairpin loop followed by four kringle domains (Fig. 1*A*). Alternate splice transcripts of *HGF* give rise to shorter protein isoforms, called HGF/NK1 and HGF/NK2, depending on the number of kringle domains encoded.

**Figure 1.**
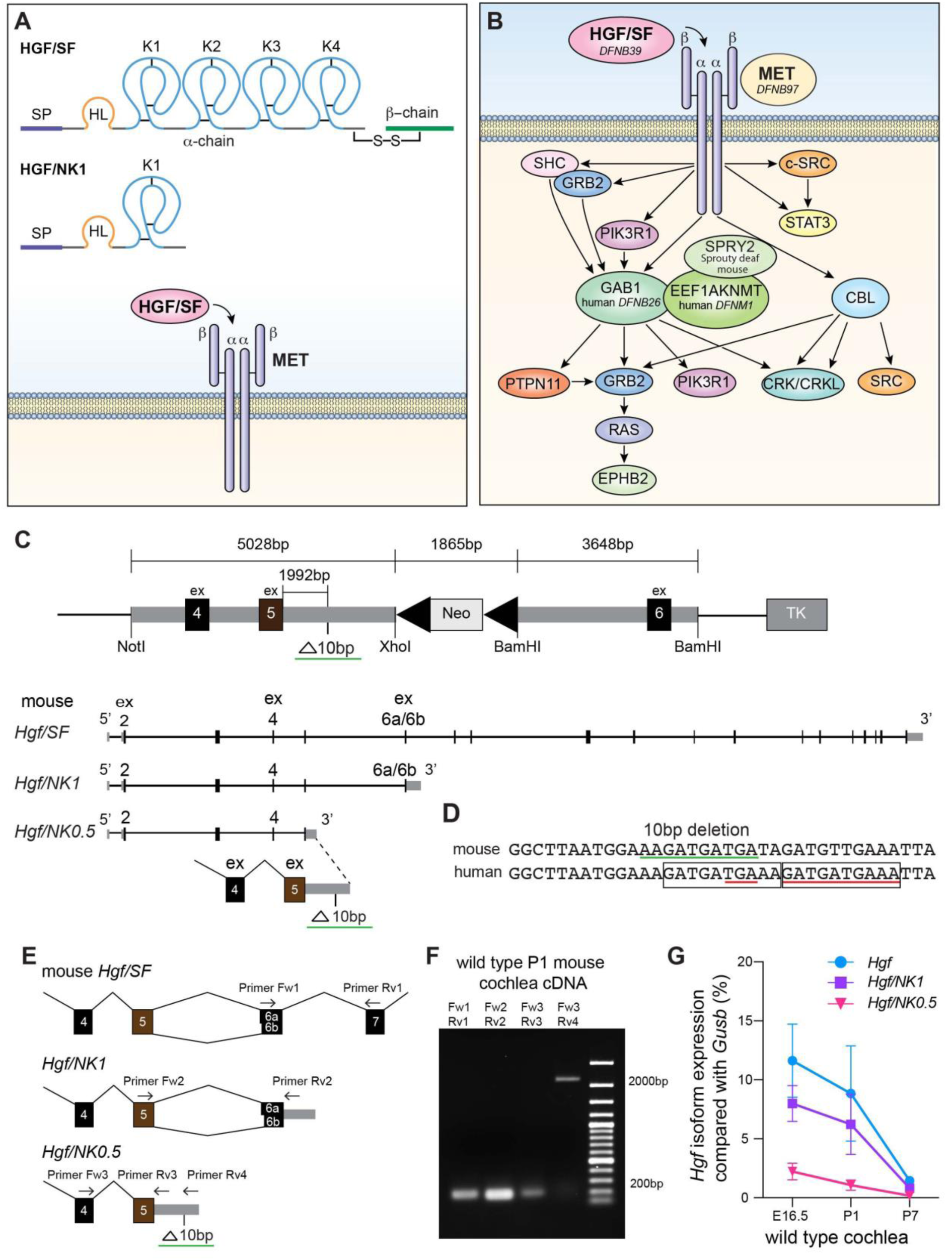
***A*,** Schematic structure of hepatocyte growth factor (HGF) which binds to and activates the MET receptor. The N-terminus signal peptide (SP) of HGF is removed by a signal peptidase. HGF is proteolytically cleaved extracellularly into a heavy alpha-chain that has a hairpin loop (HL), four kringle domains (K1-K4), each characterized by three disulfide bonds and a light beta-chain with sequence similarity to serine proteases but catalytically inoperative. ***B***. HGF-MET signaling pathways (abridged). Protein designations in upper case are the official human nomenclature (HGNC) and mouse nomenclature protein acronyms (Mouse Genome Informatics) as of August 2019. For example, the previous METTL13 is a synonym for EEF1AKNMT and SHP-2 is a synonym for PTPN11. Proteins are noted whose genes are associated with human and mouse hereditary deafness. ***C***. Targeting construct used to generate a mouse 10 base pair (bp) deletion mutation that models human deafness DFNB39. The targeting construct has a 10bp deletion engineered into intron 5 of mouse *Hgf*. The NotI, XhoI and BamHI restriction endonuclease sites were introduced into the targeting construct. The BamH1 site downstream of exon 6 is present endogenously in mouse genomic DNA. The Neomycin (Neo) cassette is flanked by loxP sites (left pointing black triangles). Depicted below the targeting construct are the alternate transcripts of the *Hgf* locus, RefSeq NM_001289458 encodes the 728 amino acid canonical longest isoform (HGF/SF), while NM_001289461 encodes an isoform using an alternate exon 6 acceptor site, and is 5 amino acids shorted (HGF_723_). A short isoform *Hgf/NK0.5* of mouse *Hgf* (AK142159.1) is orthologous to an annotated human short *HGF* isoform (ENST00000643). *Hgf/NK0.5* encodes approximately one half (35 residues) of the first of the four kringle domains (K1; 80 residues) and differs from the well-studied short HGF/NK1 *Hgf* isoform that has an entire K1 of 80 residues. The protein coding sequence of *Hgf/NK0.5* ends with sequence in exon 5 plus one intronic nucleotide to complete a conserved arginine codon, which is followed by a conserved UAA translation termination codon and regions of conserved sequence of the 3’UTR (see Extended Figure 1-1*A* for full details). ***D*,** The deleted 10bp sequence in intron 5 (green underline) is also part of 3’UTR sequence of HGF/NK0.5. The deleted 10bp is 100% identical to human sequence and to many mammals including chimp, rhesus monkey, dog and opossum. In human, a 10bp sequence is tandemly duplicated (two boxes), one copy is deleted in some subjects with DFNB39 deafness. By convention the deletion is annotated as the second copy (http://varnomen.hgvs.org/). The location of the 3bp deletion (TGA, short red underline) is the first unambiguous sequence deleted and is also a recessive variant associated with DFNB39 nonsyndromic deafness segregating in numerous human families (Schultz et al., 2009; Richard et al., 2019). ***E***, Schematic gene structures of mouse *Hgf/SF*, *Hgf/NK1* and *Hgf/NK0.5* showing the locations of RT-PCR primers. Each primer pair was designed to detect unique sequence of *Hgf/SF*, *Hgf/NK1* or *Hgf/NK0.5*. Exon 6 has two acceptor splice sites such that the encoded sequence of exons 6a and 6b differs in length by five evolutionarily conserved residues (SFLFS). ***F***, RT-PCR analysis of portions of three *Hgf* isoforms. *Hgf/SF* (180bp), *Hgf/NK1* (181bp) and *Hgf/NK0.5* (191bp and 2229bp) are all expressed in the P1 mouse cochlea. ***G,*** Developmental expression in wild type mouse cochlea using ddPCR (digital droplet) analyses. Expression levels in the cochlea of the *Hgf*/*SF*, *Hgf*/*NK1* and *Hgf*/*NK0*.5 isoforms of the gene encoding hepatocyte growth factor (*Hgf*) relative to *Gusb* expression. Levels of expression of these three isoforms decrease from E16.5 to P7. Error bars are means ± SD for three independent biological ddPCR determinations of cDNA synthesized three different times from mRNA isolated each time from four cochlea of two mice. Each ddPCR point is the mean of three technical replicates of cDNAs from mice at E16.5, P1 and P7.

**Extended Figure 1-1.**
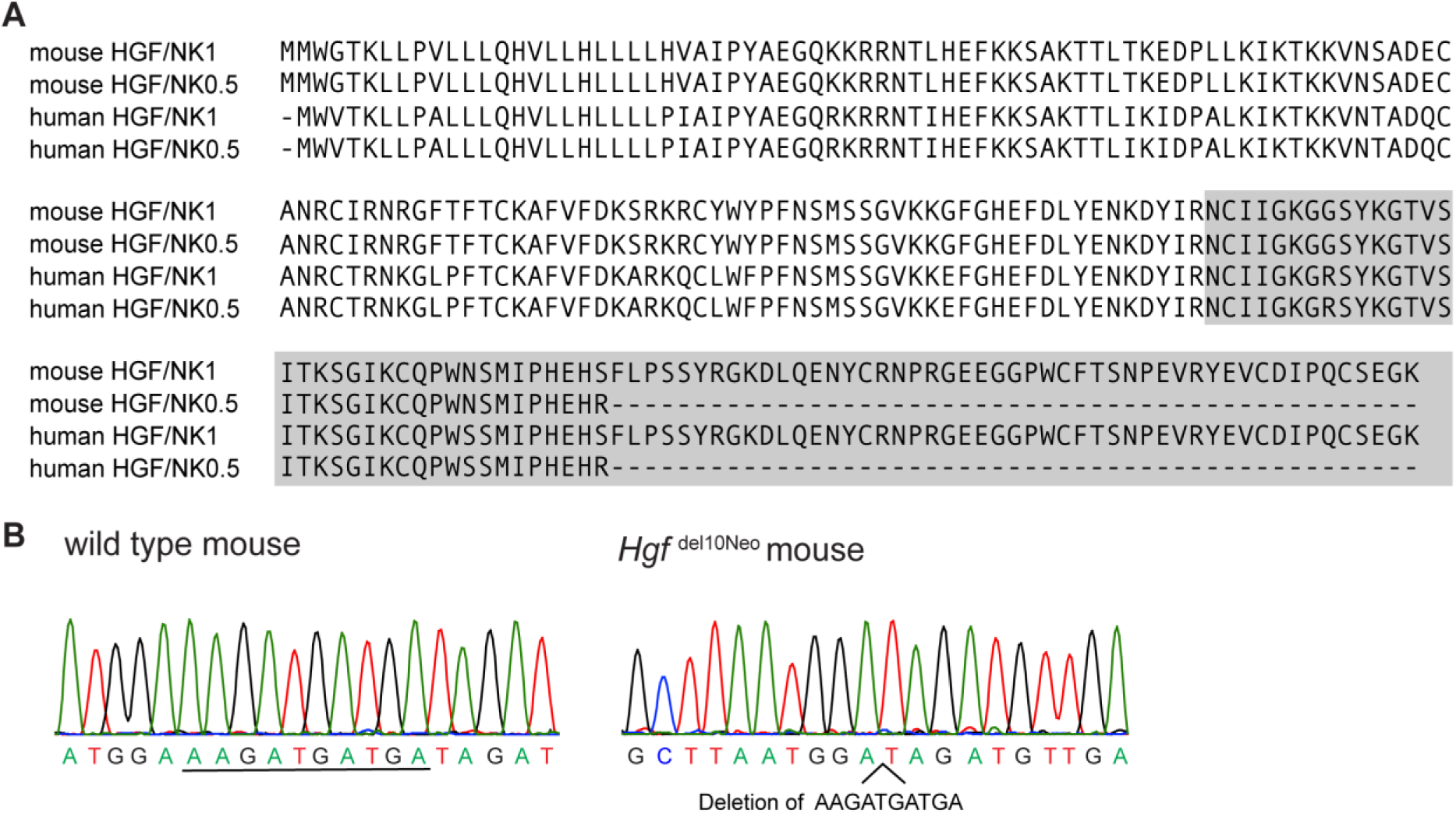
***A***, Alignment of amino acids of mouse and human HGF/NK1 and HGF/NK0.5. Kringle domain sequence is highlighted. ***B***, Representative DNA Sanger sequence traces are shown for a wild type and a Hgf del10Neo mouse.

HGF activates the MET receptor tyrosine kinase, which mediates diverse downstream pathways involved in epithelial-mesenchymal transition and the development of neural crest-derived lineages (Fig. 1*B*) (Sonnenberg et al., 1993; Birchmeier et al., 2003). When MET is active, several effector molecules are recruited that trigger signaling cascades involved in cell survival, transformation, motility and invasion, proliferation and cell cycle progression (Fig. 1*B*) as well as MET autoregulation (Organ and Tsao, 2011). HGF is also necessary for normal hearing in human and mouse. Sensorineural deafness segregating as a recessive trait in several families was genetically mapped to human chromosome 7q11.22-q21.12, and designated the *DFNB39* locus (Wajid et al., 2003). Sanger sequencing of all genes in the smallest obligate genetic linkage interval revealed 3 base pair (bp) and 10bp deletions in intron 5 of *HGF* in numerous families segregating nonsyndromic deafness (Schultz et al., 2009; Richard et al., 2019) (Fig. 1*C,D*). Interestingly, the intronic deletions occur in the 3’UTR of a previously undescribed alternative splice isoform at the *HGF* locus (referred to here as HGF/NK0.5; RefSeq NM_001010933); distinct from and smaller than HGF/NK1 (K1, Fig. 1*A*) (Cioce et al., 1996).

Homozygous knockout of mouse *Hgf* results in embryonic lethality (Kato, 2017). However, we reported that a conditional deficiency in HGF in the ear results in viable deaf mice with thinning of the stria vascularis (SV) in the cochlea (Schultz et al, 2009). Additionally, a *Hgf* transgenic mouse constitutively over-expressing HGF is also deaf, suggesting that normal cochlear development is sensitive to the amount of HGF (Takayama et al., 1996; Schultz et al., 2009). Variants of *MET* are also associated with human deafness DFNB97 (Mujtaba et al., 2015; Alabdullatif et al., 2017). In light of these observations, cochlear epithelial-specific *Hgf* and *Met* knock-out mouse models were engineered to study HGF-MET signaling during development of the SV (Shibata et al., 2016). The SV generates the +80mV endocochlear potential (EP) necessary for hair cell mechano-transduction (Wangemann, 2002). Mutations in genes expressed by SV cell types can cause a reduction or loss of EP resulting in deafness (Steel and Barkway, 1989; Tachibana, 1999). Deficits of either HGF or MET resulted in a failure of neural crest cells to incorporate into the intermediate cell layer of the SV (Shibata et al., 2016), providing an explanation for the abnormally thin SV in *Hgf* conditional knockout mice (Schultz et al., 2009).

Despite evidence related to the roles of HGF in hearing, the question remained as to whether the noncoding variants of human *HGF* described in Schultz et al. are benign but point to the actual closely-linked deafness-causing variants, referred to as linkage disequilibrium. Alternatively, are the noncoding variants the direct cause of cochlear pathology in DFNB39 deafness? If the latter, then what’s the mechanism? To answer these questions, a mouse model was engineered with a 10bp deletion in the homologous region in mouse (Fig. 1*D*) (Schultz et al., 2009). The phenotype of this mouse recapitulates human DFNB39 deafness and reveals the pathophysiological mechanism of neurosensory deafness.

## Materials and Methods

### Mouse model of DFNB39 human deafness

A targeting construct comprising 5028bp (left arm) and 3648bp (right arm) of mouse genomic DNA from the *Hgf* locus was cloned into a targeting vector based on pPNT-loxP-Neo (Fig. 1*C*). The right and left arms spanned exons 4, 5 and 6. A 10bp deletion in intron 5 was introduced by site-directed mutagenesis. In human DFNB39 families, homozygous 3bp and 10bp deletions occur in a highly conserved region of intron 5 (Schultz et al., 2009). We deleted a 10bp sequence that is 90% identical between mouse and human and overlaps the region of identity corresponding to both the 3bp (c.482+1986_1988delTGA; RefSeq NM_000601) and 10bp (c.482+1991_2000delGATGATGAAA) deletions in humans (Fig. 1*D*). In humans, this 10bp sequence is the first of two identical tandem 10bp sequences (Fig. 1*D*). A 1.8kb neomycin (neo) selection cassette was introduced into intron 5. Targeting constructs corresponding to both mutant and wild type sequences were provided to the University of Michigan Transgenic Animal Model Core for electroporation into Bruce4 ES cells. Mutant containing ES cells were recovered and two independent lines, designated 11224 and 11225, were established with the targeted mutation of *Hgf*. The 11225 line was crossed with a ZP3-cre mouse (de Vries et al., 2000) and the Neo cassette was excised, while a separate 11225 line with the Neo cassette intact was maintained. The JAX mouse nomenclature committee designated the lines B6.Cg-*Hgf*^Tm1.1Tbf^ (MGI:6294040, founder line with neo cassette) and B6.Cg-*Hgf* ^Tm1Tbf^ (MGI:6294042, neo cassette removed), referred to here as *Hgf*^del10Neo^ and *Hgf*^del10^, respectively. Back-crossing of each line to C57BL/6J continued for at least six generations before crosses between heterozygotes were performed to generate mice for auditory evaluations.

### Evaluation of B6.Cg-Hgf^Tm1.1Tbf^ mice by the NIH Phenotyping Service

Founder males for B6.Cg-*Hgf* ^Tm1.1Tbf^ (*Hgf* ^del10Neo^) were backcrossed to C57BL/6J, and heterozygous pups were crossed to generate mice for evaluation by the Mouse Phenotyping Service, Division of Veterinary Resources at the NIH. Eight wild type (WT), six heterozygous (HET) and four homozygous knock-in (KI; i.e. *Hgf* ^del10Neo/del10Neo^) mice, for a total of 18 mice, were evaluated at age 3 months. There were equal numbers of males and females of each genotype. The phenotype assessment was a comprehensive evaluation of major organ weights, hematology, serum chemistries, gross and microscopic organ evaluation.

### Generation of Hgf conditional knock-out mice

*Hgf* conditional knock-out mice were generated as previously described (Shibata et al., 2016). Briefly, *Pax2*-Cre [Research Resource Identifier (RRID): MMRRC_010569-UNC; CD1 background] and *Hgf*-floxed (RRID: MMRRC_000423-UNC; B6/129 hybrid) were utilized to generate cochlear epithelium-specific deletions of *Hgf*.

### Immunohistochemistry and measurements of strial thickness

For immunohistochemistry of cochlear sections, fixed adult mouse inner ears were decalcified in 150 mM EDTA for 5-7 days, transferred to 30% sucrose and then embedded and frozen in SCEM tissue embedding medium (C-EM001, Section-Lab Co, Ltd.; Hiroshima, Japan). Adhesive film (C-FUF303, Section-Lab Co, Ltd.) was fastened to the cut surface of the sample in order to support the section and cut slowly with a blade to obtain 10 µm thickness sections. The adhesive film with sections attached was submerged for 60 seconds in 100% ethanol, then transferred to distilled water. The adhesive film consists of a thin plastic film and an adhesive, which prevents specimen shrinkage and detachment. This methodology allows for high quality anatomic preservation of the specimen and sectioning at a thickness of 0.5 µm. Mid-modiolar sections were obtained from each cochlea where an endocochlear potential recording had been performed.

Fluorescence immunohistochemistry for known SV cell-type markers was performed as follows. Mid-modiolar sections were washed in PBS then permeabilized and blocked for 1 hour at room temperature in PBS with 0.2% Triton X-100 (PBS-T) with 10% fetal bovine serum (A3840001, ThermoFisher Scientific, Waltham, MA). Samples were then incubated in the appropriate primary antibodies in PBS-T with 10% fetal bovine serum, followed by three rinses in PBS-T and labelling with AlexaFluor-conjugated secondary antibodies (1:250, Life Technologies) in PBS-T for 1 hour at room temperature. Where indicated, 4,6-diamidino-2-phenylindole (1:10,000, Life Technologies) was included with the secondary antibodies to detect nuclei. Organs were washed in PBS three times and mounted in SlowFade Gold (S36937, Invitrogen, ThermoFisher). Specimens were imaged using a Zeiss confocal microscope. Sections were mounted with SCEM tissue embedding medium (C-EM001, Section-Lab Co, Ltd.). Primary antibodies used included rabbit anti-KCNJ10 (RRID: AB_2040120, Alomone Labs, APC-035, polyclonal, dilution 1:200), rabbit anti-CLDN11 (RRID: AB_2533259, Life Technologies, 364500, polyclonal, dilution 1:200), goat anti-SLC12A2 (RRID: AB_2188633, Santa Cruz Biotech, sc-21545, polyclonal, dilution 1:200), goat anti-KCNQ1 (RRID: AB_2131554, Santa Cruz Biotech, sc-10646, polyclonal, dilution 1:200), Phalloidin AlexaFluor 647 (RRID: AB_2620155, Invitrogen, A22287, dilution 1:250).

### *In situ hybridization* with digoxigenin-labeled antisense riboprobes

*In situ* hybridization on postnatal day 0 (P0) cochlear cross-sections was performed as previously described (Shibata et al., 2016). Briefly, P0 mice heads were fixed in 4% paraformaldehyde in PBS overnight at 4°C, sunk in 30% sucrose in PBS at 4°C, incubated in Tissue-Tek O.C.T. compound (Sakura Finetek, USA, Inc., Torrance, CA) at room temperature for 10 min, and frozen on dry ice. Sections, 14 μm thick, were cut using a Leica 3050S cryostat. Digoxigenin-labeled antisense riboprobes were synthesized using standard protocols (Stern, 1998). The following probes were used: *Hgf*, *Aldh1a2* (gift from U. Dräger, University of Massachusetts Medical School, Worcester, MA), *Cldn11* and *Dct* (gift from A. Kispert, Hannover Medical School, Hannover, Germany). The *in situ* hybridization procedure was modified from a published protocol (Henrique et al., 1995).

### In situ hybridization (smFISH) using RNAscope probes

*In situ* hybridizations were performed using RNAscope Probe-Mm-*Hgf*-No-XHs (target region: 6-2185 nucleotides, NM_001289458.1), Probe-Mm-*Hgf*-C3 (target region: 1203-2113 nucleotides, NM_001289458.1, that is equal to 1120-2030 nucleotides of NM_010427.4) (Table 1). RNAscope probes from Advanced Cell Diagnostics (ACD, Newark, CA) were used with sections of cochleae from C57BL/6J wild type mice at embryonic ages E14.5, E18.5 and P30. Embryonic cochleae with the brain hemisected were fixed overnight at 4°C in 4% PFA in 1x PBS. Cochleae were then cryopreserved overnight in 15% and then overnight in 30% sucrose at 4°C. Adult cochleae were dissected from the head and fixed overnight at 4°C in 4% PFA in 1x PBS. Fixed adult mouse inner ears were decalcified in 150 mM EDTA for 5-7 days, transferred to 30% sucrose, and then embedded and frozen in SCEM tissue embedding medium (Section-Lab Co, Ltd.). Adhesive film (Section-Lab Co, Ltd.; Hiroshima, Japan) was fastened to the cut surface of the sample in order to support the section and cut slowly with a blade to obtain thin mid-modiolar sections. The adhesive film with section attached was submerged in 100% EtOH for 60 seconds, then transferred to distilled water. This methodology allows for high quality anatomic preservation of the specimen. Frozen tissues were sectioned (10 µm thickness) with a CM3050S cryostat microtome (Leica, Vienna, Austria). Sections were mounted with SCMM mounting medium (Section-Lab, Hiroshima, Japan) and imaged using a 1.4 N.A. objective.

### Stria vascularis measurements and fluorescence intensity quantifications of strial cell type markers

ImageJ was utilized to calculate the cross-sectional area and thickness of the SV in mid-modiolar sections of both wild type and homozygous KI mice in both the *Hgf ^del10Neo^* and *Hgf ^del10^* mouse lines at postnatal day 60 (P60). Fluorescence intensity quantification was performed in ImageJ by calculating the fluorescence intensity of the outlined region of the SV. Fluorescence intensity was normalized by comparing the SV fluorescence intensity to that of a corresponding region in the scala media. Measurements for the upper (apical), middle, and lower (basal) turns of the cochlea were obtained. These measurements were obtained for known SV cell types including intermediate cells (KCNJ10), marginal cells (SLC12A2, KCNQ1), and basal cells (CLDN11). KCNJ10 fluorescence intensity in the spiral ganglion neurons served as a control for immunofluorescence signal intensity measurements. Spiral ganglion fluorescence intensity was unchanged between KI and WT mice with no statistically significant difference between fluorescence intensity measurements (data not shown).

### Auditory Testing

Auditory Brainstem Responses (ABRs) were measured at ages 4 weeks, 8 weeks and 25 weeks after birth. Mice were anesthetized by an intraperitoneal (IP) injection of ketamine (56 mg/kg) and dexdomitor (0.375 mg/kg) and placed on a heating pad connected to a temperature controller (World Precision Instruments T-1000 or T-2000, Sarasota, FL) inside a sound-treated booth (Acoustic Systems, Austin, TX). A rectal probe was used to monitor body temperature and a heating pad was used to maintain body temperature near 37°C. Auditory brainstem responses (ABRs) were obtained using Tucker-Davis Technologies (TDT; Alachua, FL) hardware (RZ6 Processor) and software (BioSigRZ, v. 5.1; RRID: SCR_014820).

For ABR testing, subdermal needle electrodes (Rhythmlink, Columbia, SC, USA) were placed at the vertex, under the test ear, and under the contralateral ear (ground). Blackman-gated tone burst stimuli (3 msec, 29.9/sec, alternating polarity) were presented to the test ear at 8, 16, 32, and 40 kHz via a closed-field TDT MF-1 speaker. Responses were amplified (20x), filtered (.3-3 kHz) and digitized (25 kHz) with 512-1024 artifact-free responses per waveform. For each frequency, testing began at 80 dB SPL and decreased in 10 dB steps until the ABR waveform was no longer discernable. If no response was obtained at 80 dB SPL, testing was performed at a maximum level of 90 dB SPL. Once the response was lost, testing continued in 5 dB steps with a minimum of two waveforms per stimulus level to verify repeatability of ABR waves. ABR thresholds were determined by visual inspection of stacked waveforms for the lowest stimulus level that yielded repeatable waves.

### EP measurements

Methods for endocochlear potential (EP) measurement have been described (Wangemann et al., 2004; Wangemann et al., 2007). Here, mice were anesthetized with 2,2,2-tribromoethanol (T4842, Sigma-Aldrich, St. Louis, MO) at a dose of 0.35 mg/g body weight. EP measurements were made using glass microelectrodes inserted into the round window and through the basilar membrane of the first turn of the cochlea. Induction of anoxia, allowing measurement of anoxic-state EP, was accomplished by intramuscular injection of succinylcholine chloride (0.1 µg/g, NDC-0409-6629-02, Pfizer, NY, NY) after establishment of deep anesthesia followed by additional injection of 2,2,2-Tribromoethanol (T4842, Sigma-Aldrich, St. Louis, MO). Anoxic-state EP provides an indicator of the lowest EP and sensory hair cell function. In the presence of functional hair cells, the anoxic-state EP is negative, whereas the EP is zero if the hair cells are not functional. Data were recorded digitally (Digidata 1440A and AxoScope 10; Axon Instruments) and analyzed using Clampfit10 (RRID: SCR_011323, Molecular Devices, San Jose, CA).

### Transcriptomics

KI and WT littermate mice at 5-27 weeks of age were euthanized via CO_2_ asphyxiation, and dissections of the cochlea, lung, and kidney were immediately frozen in liquid nitrogen. Tissue was pulverized with a Covaris CPO2 CryoPrep Automated Dry Pulverizer (Covaris, Woburn, MA) and RNA was extracted with Trizol (Invitrogen). For RNA-seq, total RNA from cochleae was reverse transcribed with random primers after ribo-depletion using a TruSeq library kit (15031048 Rev. C, Illumina, San Diego, CA), and 2×93bp sequenced on an Illumina HiSeq1500 instrument. The reads were mapped to the mouse genome (GRCm38.vM11) using STAR (Dobin et al., 2013). Differentially expressed (DE) genes were determined by DeSeq2 (Love et al., 2014). A total of 19 DE genes were found to be downregulated, and 14 DE genes upregulated, in *Hgf* ^del10Neo/del10Neo^ cochleae (GEO Accession ID: GSE137721). The list of DE genes was analyzed by EnrichR for gene ontology analyses (RRID:SCR_001575; http://amp.pharm.mssm.edu/Enrichr/) as previously described (Chen et al., 2013; Kuleshov et al., 2016; Pazhouhandeh et al., 2017). Enrichr is an integrated web-based application that includes updated gene-set libraries, alternative approaches to ranking enriched terms, and a variety of interactive visualization approaches to display the enrichment results. Enrichr employs three approaches to compute enrichment as previously described (Jagannathan et al., 2017). The combined score approach where enrichment was calculated from the combination of the p-value computed using the Fisher exact test and the z-score was utilized. RT-PCR was performed using C57BL/6J wild type P1 mouse cochlea cDNA and PCR products were Sanger sequenced. Quantitative RT-PCR were performed using ddPCR (digital droplet) Supermix for Probes (186-3024, BioRad, Hercules, CA) on a Bio-Rad QX200 droplet digital PCR system (BioRad). Each experiment was repeated three times and expression levels were calculated as ratio of positive *Hgf* cDNA droplets to droplet positive for the housekeeping gene *Gusb*.

qRT-PCR was performed using TaqMan Gene Expression Master Mix (4369514, ThermoFisher) on a ViiA7 Realtime PCR instrument (Applied Biosystems). Taqman probe for *Hgf* spanned the exon 3-exon 4 junction, which should recognize all isoforms of *Hgf* or were specific to the exon 5-exon 6a and exon 5-exon 6b junctions. Expression levels of *Hgf* probe 3-4 were calculated as the delta Cq between the target probe and the geometric mean of reference probes *Actb* and *Gusb*. The 6a versus 6b alternative splice acceptor site usage was calculated as the Log2 fold difference for each sample. Details for primers and Taqman probes utilized are listed in Table 2.

### Statistical Analyses

For pairwise comparisons between mutants and wild type littermates, an unpaired 2-tailed Student’s t-test was used. ANOVA was used for comparisons between genotypes involving multiple tissues. Differences between genotypes within tissues were calculated using Sidak’s multiple comparisons test. All statistical analyses were performed using GraphPad Prism version 6.0 (RRID: SCR_002798; GraphPad) for PC. For measurements of strial thickness, cross-sectional area and fluorescent intensity, all values are means ± standard deviations (SD). Both males and females were tested, and there was no evidence of a significant effect of sex on any measures. Therefore, the data displayed in all graphs are from males and females combined.

## Results

### Overall phenotype of 10bp deletion homozygotes

Wild type (WT), heterozygous (HET) and homozygous knockin (KI; *Hgf* ^del10Neo/del10Neo^) mice were evaluated at age P90 (3 months) by the Mouse Phenotyping service at the NIH Division of Veterinary Resources. This comprises a comprehensive analysis of gross and histopathologic evaluations including organ weights, serum chemistries, hematology and histology. There were no obvious differences between genotypes in body size, weight or coat pigmentation. Gross inspection of the major organs and histopathology of organs, including those that normally show high expression of *Hgf* (lung, liver, kidney), showed no abnormalities in *Hgf* ^del10Neo/del10Neo^ mice. A schematic of the organ of Corti and the stria vascularis which reside in the cochlea are shown (Fig. 2*A*). The organ of Corti is composed of one row of inner hair cells and three rows of outer hair cells surrounded by supporting cells. The stria vascularis is composed of three cellular layers, which consist predominantly of marginal, intermediate and basal cells, respectively. Despite the lack of abnormalities in other organs, the inner ears of *Hgf* ^del10Neo^ homozygous KI mice exhibited gross defects including thin and detached SV, missing or pyknotic hair cells and supporting cells along with general atrophy of the organ of Corti and degeneration of the spiral ligament, among other malformations (Fig. 2*B,C*). These defects occurred in an otherwise normally developed and patterned cochlea. The organ of Corti developed with apparently all cell types present and morphologically identifiable, although reduced in numbers or showing signs of degeneration. Circling behavior and head-bobbing were not noticed in KI mice. No significant differences attributable to genotype were found in any other organ system.

**Figure 2.**
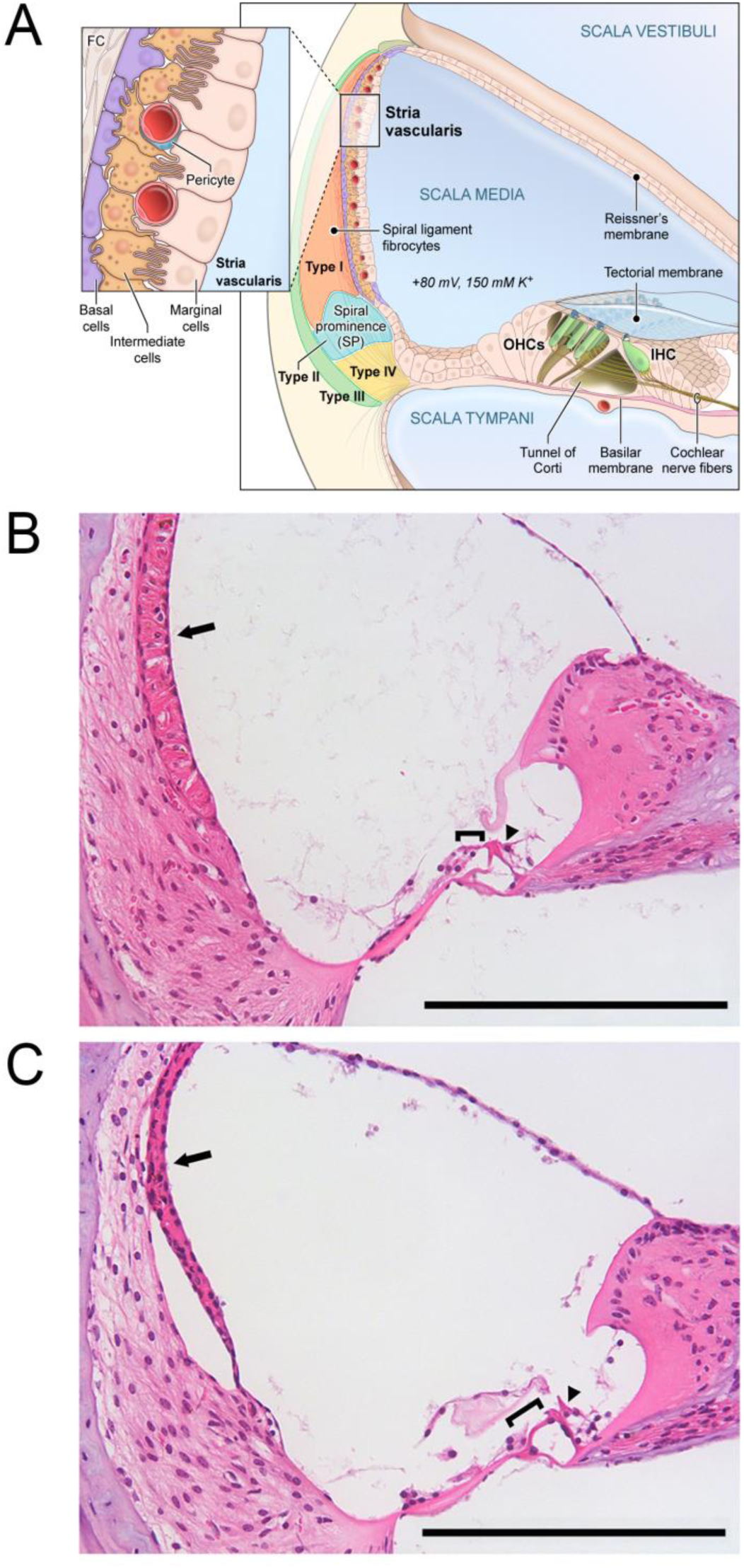
***A***, Schematic of the adult cochlea including the organ of Corti and the stria vascularis. The organ of Corti is housed within the endolymph-containing scala media and is composed of inner and outer hair cells surrounded by supporting cells. The stria vascularis is composed of three cellular layers consisting of marginal, intermediate, and basal cells. The stria vascularis generates the +80 millivolt (mV) endocochlear potential (EP). Inner hair cell (IHC), outer hair cell (OHC), fibrocyte (FC), spiral prominence (SP). ***B-C***, Hematoxylin and eosin (H&E) staining of adult P90 homozygous KI mice demonstrate changes in cochlear structure including loss of outer hair cells, strial atrophy and spiral ligament atrophy. ***B***, Representative H&E stained section of P90 WT littermate from *Hgf* ^del10Neo^ mouse line. Structures denoted as follows: stria vascularis (arrow), inner hair cell (arrowhead), outer hair cells (bracket). ***C***, Representative H&E stained section of P90 homozygous KI mouse *Hgf* ^del10Neo^ mouse line. Note detachment and thinning of the stria vascularis (arrow) from spiral ligament, spiral ligament atrophy and loss of outer hair cells (bracket) in the *Hgf* ^del10Neo^ homozygous KI mouse. Scale bar is 200 μm. Arrowhead (inner hair cell), Bracket marks outer hair cell region. Arrow points to the stria vascularis. Similar pathology was noted between *Hgf* ^del10Neo^ and *Hgf*^del10^ KI mouse lines at P90.

### Homozygous KI mice exhibit hearing loss

Homozygous KI mice from both mouse lines (*Hgf* ^del10Neo^ and *Hgf* ^del10^) were characterized by determining hearing thresholds by measuring Auditory Brainstem Responses (ABR) to pure-tone stimuli. Homozygous *Hgf* ^del10Neo^ KI mice displayed profound hearing loss at 4 weeks of age (Fig. 3*A*), which was unchanged at 8 weeks and 25 weeks. No significant differences in ABR thresholds were seen between heterozygotes or wild type littermates. In the *Hgf* ^del10^ mouse line, homozygous KI mice had moderate-to-severe hearing loss at age 4 weeks (Fig. 3*A*) that was unchanged at 8 and 25 weeks. ABR thresholds were more variable in the *Hgf* ^del10^ versus the *Hgf* ^del10Neo^ homozygotes. *Hgf* ^del10^ mice never displayed the profound hearing loss observed for *Hgf*^del10Neo^ homozygotes.

**Figure 3.**
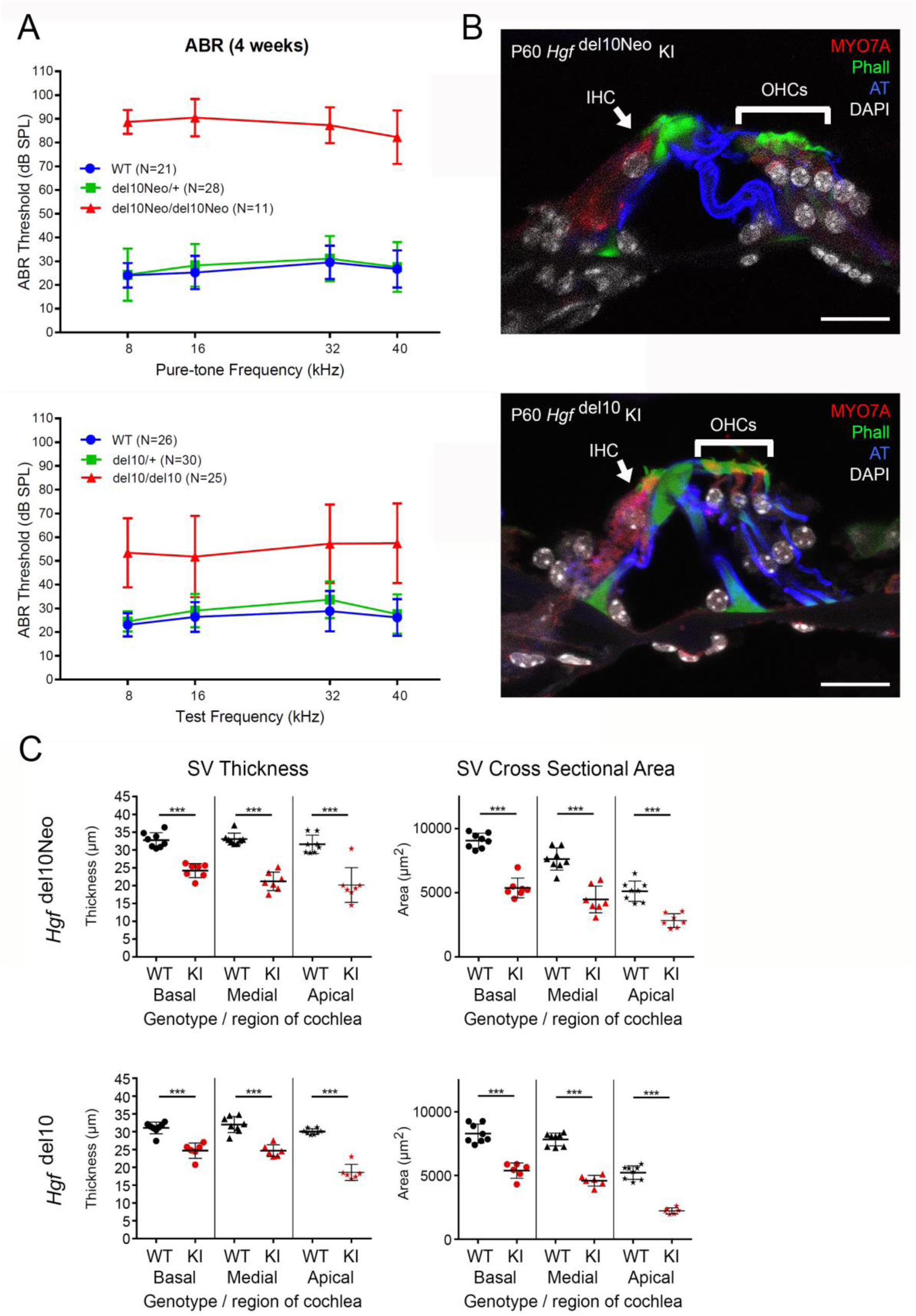
Hearing loss phenotyping and cochlear structural observations in *Hgf* ^del10Neo/del10Neo^ mice and *Hgf* ^del10/del10^ mice at P60. ***A*,** Auditory brainstem response (ABR) for *Hgf* ^del10Neo^ KI (upper panel) and *Hgf* ^del10^ KI (lower panel) mouse lines at 4 weeks. Tone burst stimuli were delivered and responses were measured at 8, 16, 32, and 40 kHz as described in the methods. ***B*,** Representative midmodiolar cross-sections of P60 organ of Corti from *Hgf* ^del10Neo/del10Neo^ mice (upper panel) and *Hgf* ^del10/del10^ mice (lower panel). Sections are stained with MYO7A for hair cells (red), Phalloidin (green), acetylated tubulin for supporting cells (AT, blue), and DAPI for nuclei (white). Note inner hair cells and outer hair cells in red labeled with anti-MYO7A antibody. Scale bars are 20 μm. Inner hair cell (IHC) is denoted with label and arrow. Outer hair cells (OHCs) are denoted by a label and bracket. ***C*,** Strial thickness and strial cross-sectional area measurements in both *Hgf* ^del10Neo^ (upper panels) and *Hgf* ^del10^ (lower panels) homozygous KI mice compared to wild type (WT) littermate controls.

### Histological evaluation of homozygous KI mice indicate that hearing loss originates from defects in the stria vascularis

For both *Hgf* ^del10Neo^ and *Hgf* ^del10^ KI lines at P60, examination of the organ of Corti with fluorescent immunohistochemistry demonstrated a normal morphology and complement of inner and outer hair cells and supporting cells (Fig. 3*B*), but at P90 there was a loss of outer hair cells (Fig. 2*A,B*). The *Hgf* ^del10Neo/del10Neo^ mice exhibited a profound hearing loss starting at 4 weeks of age. In contrast, strial thinning (Fig. 3) and reduced KCNJ10 expression was observed at P60 (Fig. 4) that preceded degenerative changes in the organ of Corti (Fig. 2*A,B*). After a gross wild type developmental patterning of the cochlea, we sought to identify the primary defect causing the hearing loss in the homozygous *Hgf* ^del10Neo^ and *Hgf* ^del10^ KI lines.

**Figure 4.**
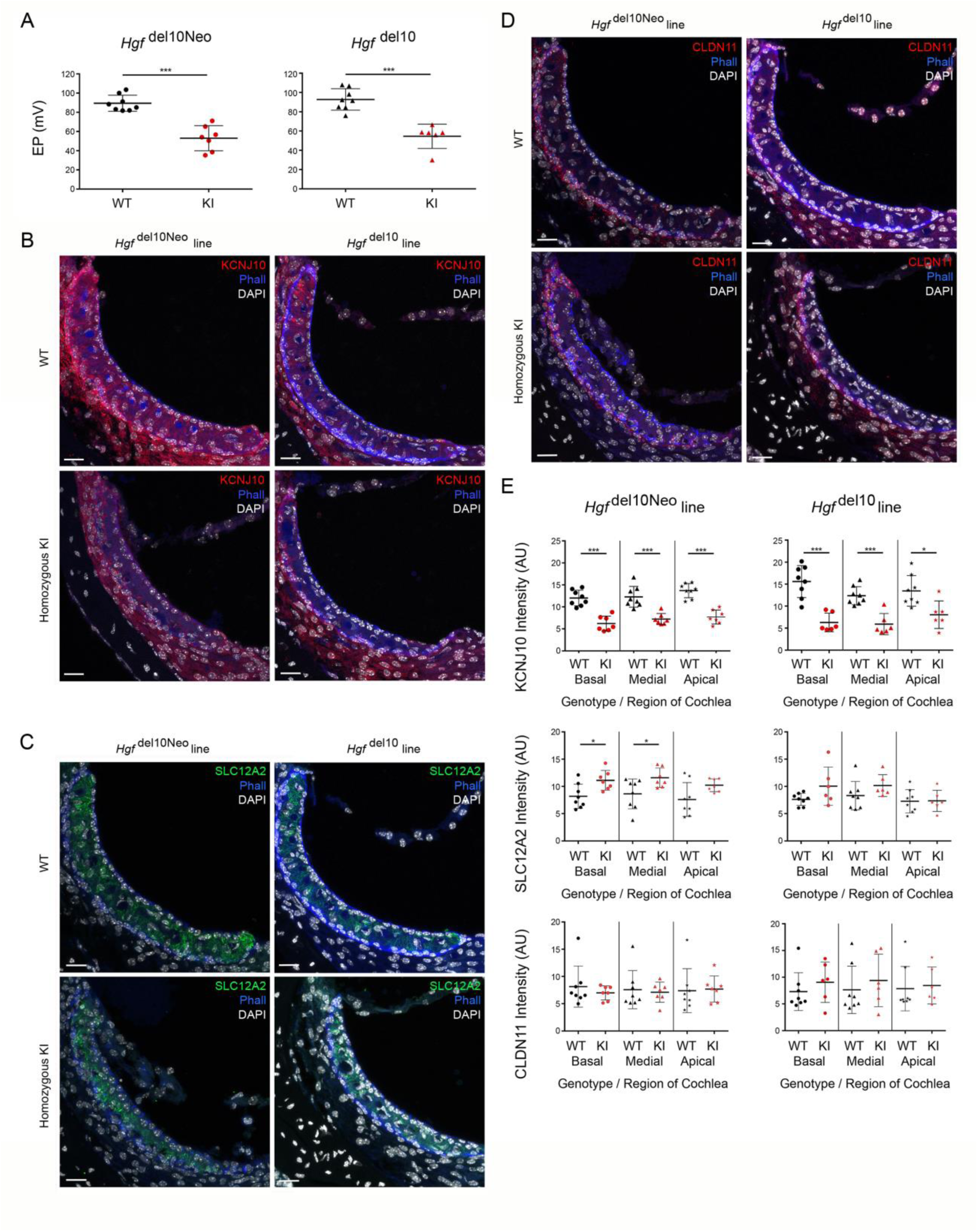
Physiological and histological evaluation of the *Hgf* ^del10Neo^ and *Hgf* ^del10^ lines demonstrate deficits in stria vascularis melanocytes. ***A*,** Endocochlear potential (EP) measurements in the *Hgf* ^del10Neo^ (left panel) and *Hgf* ^del10^ (right panel) mouse lines demonstrate significant differences between homozygous KI and wild type littermates (unpaired 2-tailed Student t-test, p < 0.001). ***B*,** Representative immunostaining for KCNJ10 in the *Hgf* ^del10Neo^ and *Hgf* ^del10^ lines demonstrate visually apparent decrease in KCNJ10 immunostaining in the homozygous KI mice from both mouse lines. WT mice are shown in the top two panels and homozygous KI mice are shown in the bottom two panels. Sections are stained with KCNJ10 for intermediate cells (red), Phalloidin for tight junctions (blue), and DAPI for nuclei (white). ***C*,** Representative immunostaining for SLC12A2 in the *Hgf* ^del10Neo^ and *Hgf* ^del10^ lines demonstrate SLC12A2 immunostaining in the homozygous KI mice from both mouse lines. Differences in immunostaining are not readily apparent. WT mice are shown in the top two panels and homozygous KI mice are shown in the bottom two panels. Sections are stained with SLC12A2 for marginal cells (red), Phalloidin for tight junctions (blue), and DAPI for nuclei (white). ***D*,** Representative immunostaining for CLDN11 in the *Hgf* ^del10Neo^ and *Hgf* ^del10^ lines demonstrate CLDN11 immunostaining in the homozygous KI mice from both mouse lines. No differences are apparent in CLDN11 immunostaining. WT mice are shown in the top two panels and homozygous KI mice are shown in the bottom two panels. Sections are stained with CLDN11 for basal cells (red), Phalloidin for tight junctions (blue), and DAPI for nuclei (white). ***E***, Quantitative intensity analysis of KCNJ10, SLC12A2, and CLDN11 immunostaining in the *Hgf* ^del10Neo^ and *Hgf* ^del10^ lines (n ≥ 6 animals per line). A significant decrease in KCNJ10 intensity in homozygous KI mice is shown for both mouse lines. For SLC12A2 immunostaining, only the *Hgf* ^del10Neo^ mouse line demonstrates significant increase in SLC12A2 intensity in homozygous KI mice, while no significant change is apparent between homozygous KI and WT mice in the *Hgf* ^del10^ mouse line. Finally, no significant change in CLDN11 intensity is seen in either mouse line.

The thinning and sometimes detachment of the SV seen in the KI mice was reported in a mouse with a conditional deletion of *Hgf* exon 6 and in a constitutively overexpressing *Hgf* mouse (MH19) (Takayama et al., 1996; Schultz et al., 2009). This was especially interesting given that expression of *Hgf* and *Met*, encoding the HGF receptor tyrosine kinase, is expressed in the early developing SV, and HGF–MET signaling has been shown to be crucial to the incorporation of melanocytes into the SV (Shibata et al., 2016). Therefore, we made detailed measurements of strial thickness and total area at three defined locations in the cochlea, comparing P60 WT to *Hgf*^del10Neo^ and *Hgf* ^del10^ lines (Fig. 2*E*). Significant reductions in both the *Hgf* ^del10Neo^ and *Hgf*^del10^ lines compared to WT in thickness and total strial area were noted at the base, mid-turn and apex of the cochlea (unpaired 2-tailed Student t-test, p < 0.001).

Given the possibility of a SV defect as the primary site of pathology associated with hearing loss, and the relationship between SV function and ion homeostasis in the inner ear (Hibino et al., 2010; Patuzzi, 2011), we measured endocochlear potentials (EPs) in P60 mice as a direct measure of SV function. The EP has been shown to be directly proportional to strial volume (Schulte and Schmiedt, 1992). Homozygous *Hgf* ^del10Neo^ and *Hgf* ^del10^ lines showed statistically significant reduction in their EPs when compared to their WT littermates (Fig. 4*A*; unpaired 2-tailed Student t-tests, p < 0.001).

The SV is composed of three cellular layers: marginal, intermediate and basal cells (Fig. 2*A*). Examination of the SV with immunohistochemistry using antibodies specific for each layer (Fig. 4*B-D*) indicates that intermediate cells are deficient, as indicated by the expression of the intermediate cell specific marker, KCNJ10, in homozygous KI mice compared to WT mice (Fig. 4*B,E*) (p < 0.0001). However, notably KCNJ10 immunostaining is not absent in homozygous KI mice. By comparison, markers for the other two strial cell types are not reduced in homozygous KI mice (Fig. 4*C,D*). Immuno-staining intensities for SLC12A2, a marker for marginal cells (Fig. 4*C,E*), are slightly increased in the apical and basal turns of the *Hgf* ^del10Neo^ homozygous KI mice compared to the SLC12A2 intensities in WT littermates while they are unchanged in the *Hgf* ^del10^ homozygous KI mice (p < 0.01) (Fig. 4*C,E*). Expression intensities for CLDN11, a tight junction protein and marker for basal cells, are indistinguishable for KI and WT littermates from both mouse lines (Fig. 4*D,E*). This finding was confirmed and extended by *in situ* hybridization experiments with probes for *Cldn11* (basal cells), *Aldh1a2* (marginal cells) and *Dct* (intermediate cells) (Fig. 5*A-L*) in *Hgf*-conditional knock out (CKO) mice at P0. The expression of *Aldh1a2* and *Cldn11* is unchanged between WT, heterozygotes, and homozygous *Hgf*-CKO mice. The images for *Dct* are particularly illuminating since this gene encodes dopachrome tautomerase, which is expressed by melanocytes; the cells that constitute the future SV intermediate cell layer. The *Dct* signal is markedly reduced, but not absent, at P0 in *Hgf*-CKO mice compared to WT (Figure 5*A-F*). These data are consistent with previous observations (Shibata et al., 2016) and reminiscent of the reduced KCNJ10 immunostaining seen in homozygous KI (*Hgf* ^del10/del10^) mice in our mouse model. Taken together, the data implies that the primary defect in *Hgf* ^del10/del10^ mice is a significant reduction in the number of neural crest derived melanocytes that infiltrate the developing SV.

**Figure 5.**
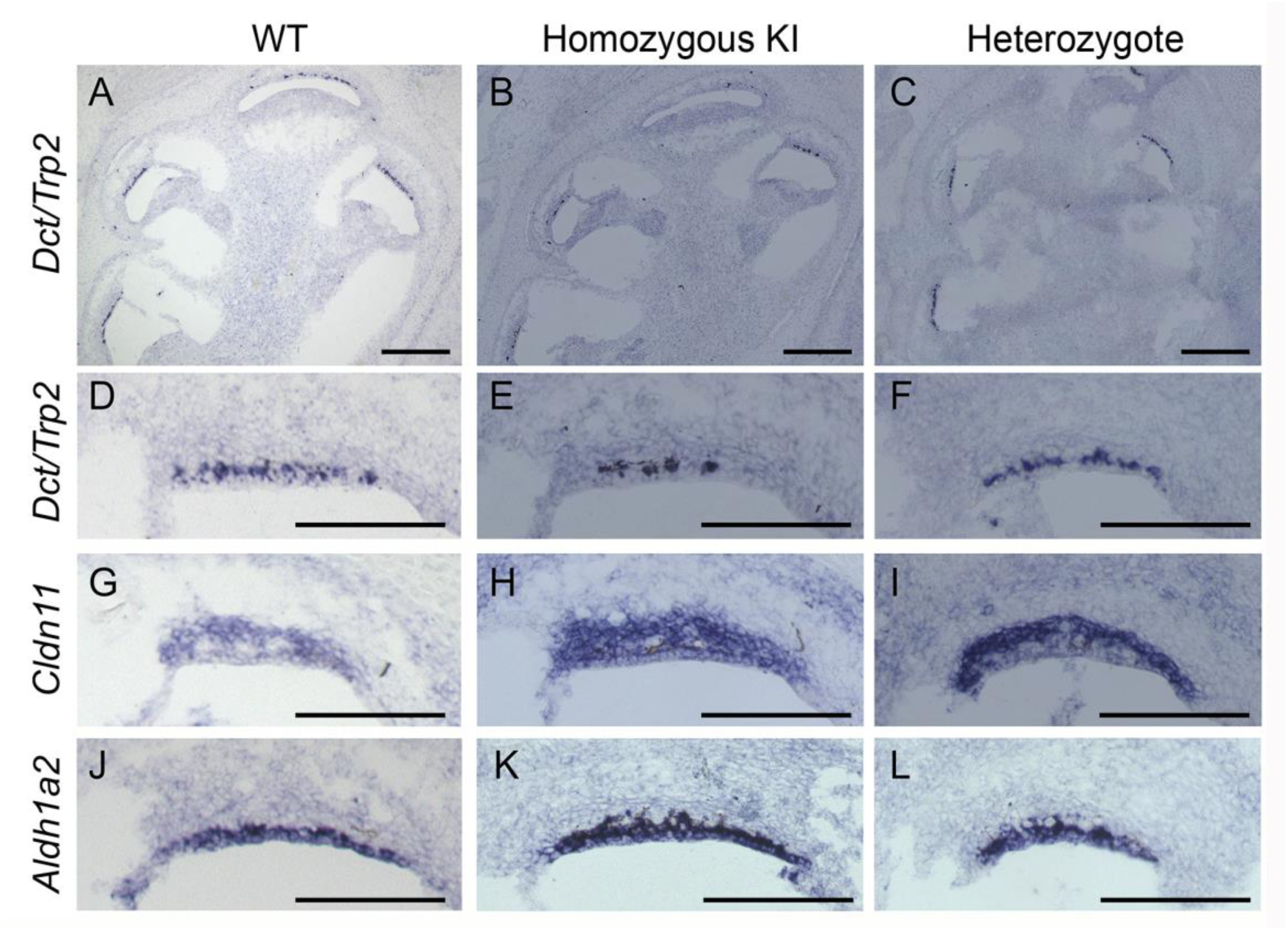
Observations in *Hgf*-CKO mice at P0 confirm a reduction in future strial intermediate cells. ***A-F*,** *In situ* hybridization for *Dct* demonstrates a reduction of *Dct* signal in homozygous CKO mice versus WT and homozygous CKO mice. Low magnification imaging of Dct signal in mid-modiolar cross sections of wild type (***A***), homozygous (***B***) and heterozygous (***C***) cochleae from *Hgf*-CKO mice. All scale bars are 100 μm. High magnification of representative stria vascularis from wild type (***D***), homozygous (***E***), and heterozygous (***F***) *Hgf*-CKO mice. *Dct* signal identifies intermediate cell layer. ***G-I*,** *In situ* hybridization for *Cldn11* demonstrates unchanged signal in wild type (***G***), homozygous (***H***), and heterozygous (***I***) *Hgf*-CKO mice. *Cldn11* signal identifies basal cell layer. ***J-L*,** *In situ* hybridization for *Aldh1a2* demonstrates unchanged signal in wild type (***J***), homozygous (***K***), and heterozygous (***L***) *Hgf*-CKO mice. All scale bars are 100 μm. *Aldh1a2* signal identifies marginal cell layer.

### The del10 intronic mutation results in altered *Hgf* expression in the inner ear

We evaluated *Hgf* mRNA levels in the cochleae, kidneys and lungs of KI mice and their WT littermates using Taqman assays and qRT-PCR. Using Taqman assays that span the constitutively expressed exons 3 and 4 (probe 3-4), which recognize all known splice isoforms, the expression level of *Hgf* was approximately 70% lower in cochlea of KI mice compared to WT (p = 0.0004, p = 0.03, respectively) in both *Hgf* ^del10Neo^ and *Hgf*^del10^ lines (Fig. 6*A,B*). In kidney and lung, there were no expression differences between WT and KI mice (p = 0.11 and 0.79 for kidney, and p = 0.64 and 0.99 for lung). Probes specific to the junction using the exon 6a or exon 6b alternative splice acceptor sites were used to evaluate the relative ratio of 6a to 6b isoform usage among *Hgf* transcripts. The 6a/6b ratios were reduced in KI compared to WT mice in both lines (p = 0.03 and p = 0.01, respectively) and these changes were not observed in the kidney and lung (all p-values > 0.03 for kidney and lung, both lines) (Fig. 6*B*). These data suggest that the noncoding 10bp deletion in *Hgf* intron 5 alters *Hgf* expression perhaps exclusively in the cochlea.

**Figure 6.**
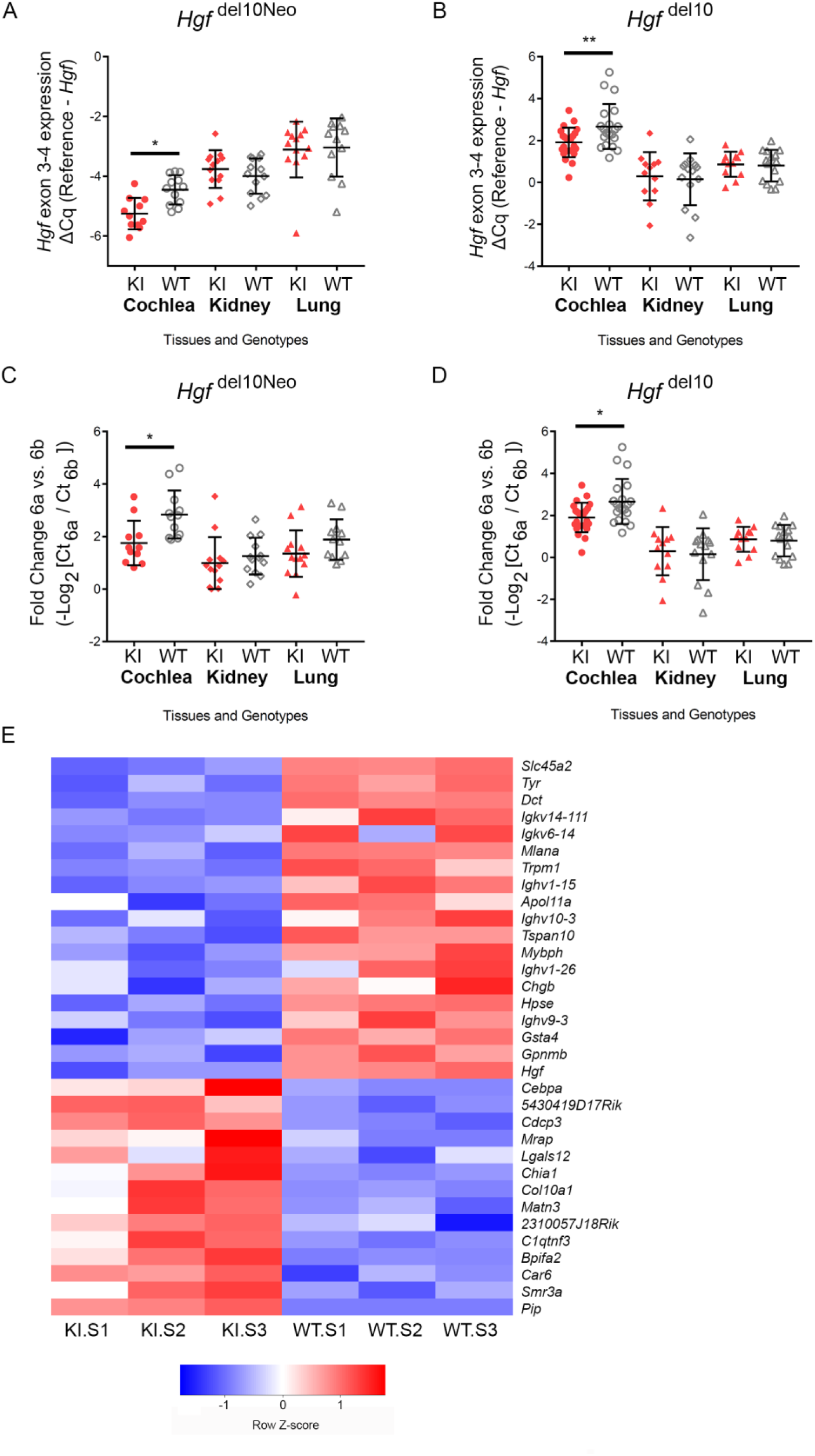
The del10 intronic *Hgf* mutation results in an inner ear alteration in *Hgf* expression. ***A-B***, Cochlear *Hgf* expression as determined by qRT-PCR using Taqman probe that spans exon 3-exon4 (detects all isoforms) is significantly lower in KI compared to WT mice in both the *Hgf* ^del10Neo^ (***A***) and *Hgf* ^del10^ (***B***) lines (p = 0.0004, p = 0.03, respectively). Graph depicts relative quantification of Hgf probe 3-4 versus the geometric mean of two reference genes (Actb and Gusb). More negative numbers indicate larger target probe Cq relative to stable reference gene Cq, and thus lower expression levels. Expression of *Hgf* in the mouse kidney and lungs is unchanged KI compared to WT mice (p = 0.11 and 0.79 for kidney, and p = 0.64 and 0.99 for lung). Sidak’s multiple comparisons test utilized to determine significance between KI and WT values after one-way ANOVA. * p < 0.05, ** p < 0.001. ***C-D***, Use of Taqman probes specific for exon 6a or exon 6b splice acceptor sites demonstrate significant but slight change in the relative ratio of 6a versus 6b alternative splice acceptor usage between WT and KI mice in the cochlea in both the *Hgf* ^del10Neo^ (***C***) and *Hgf* ^del10^ (***D***) lines (p = 0.03 and p = 0.01, respectively) while no change in this ratio was noted in the kidney and lung (all p-values > 0.03 for kidney and lung, both lines). ***E,*** RNA-Seq comparing transcriptome profiles of whole cochlea from *Hgf* ^del10Neo^ homozygous KI (KI.S1, KI.S2, KI.S3) to transcriptome profiles of whole cochlea from WT littermates (WT.S1, WT.S2, WT.S3) demonstrates downregulation of genes consistent with a loss of SV intermediate cells. Heatmap illustrates the 33 top differentially expressed genes (DEGs) including 19 genes that were down-regulated and 14 genes that were upregulated in the KI transcriptomes. Higher red intensity corresponds to higher relative expression; higher blue intensity corresponds to lower relative expression. Histogram color bar is shown for reference. WT (wild type), KI (knock-in).

Differential gene expression analysis revealed 34 significantly differentially expressed genes (DEGs) of which 19 genes were upregulated and 14 genes were downregulated (Fig. 6*C*). Among the few downregulated DEGs are two genes, *Dct* and *Tyr*, that are specific to the intermediate cells in the SV and two genes, *Slc45a2* and *Mlana*, that are known to be expressed by melanocytes. Analysis with Enrichr reveals that the downregulated genes are involved in *Mitf* signaling (TRRUST Transcription Factors 2019, Transcription Factors PPI). Gene ontology (GO) analysis (GO Biological Process, GO Molecular Function, GO Cellular Component) reveals that these genes, such as tyrosinase, are involved in the melanin synthesis pathway (Fujita et al, 2017). Melanin may have a protective role, possibly as a ROS scavenger (Bustamante et al., 1993). These RNA-Seq analyses provide independent confirmation of a loss of intermediate cells in the setting of the *Hgf del10* mutation.

The preferential reduction of *Hgf* in the cochlea in the case of the del10 mutation coupled with the confinement of differentially expressed genes to melanocyte-specific genes and pathways also suggest that the effect of the *Hgf* del10 mutation may be confined to the cochlea. Thus, this mutation appears to represent a unique situation, since neurocristopathies most often result in syndromes involving multiple organ systems (Bolande, 1974; Vega-Lopez et al., 2018; Ritter and Martin, 2019).

### Hgf RNA expression indicates a critical role in the developing stria vascularis

Single molecule fluorescent *in situ* hybridization (smFISH) using two RNAscope probes was used to detect *Hgf* transcript expression in the SV during development and adulthood. One probe (Mm-*Hgf*-No-XHs) detected the full-length sequence of mouse *Hgf* and the other probe (Mm-*Hgf*-C3) detected the *Hgf* RNA sequence towards the C-terminus (Fig. 7*A*). Both probes detected the full-length *Hgf* mRNA in the SV of E14.5, E18.5 and P30 wild type mice, providing dual confirmation of *Hgf* mRNA in the SV (Fig. 7*B,C*). smFISH labeling demonstrates the presence of both Mm-*Hgf*-No-XHs (red) and Mm-*Hgf*-C3 (blue) in marginal cells at E18.5 wild type mice (Fig. 7*B*). In adult wild type mice, *Hgf* mRNA expression is present as measured by both smFISH probes, Mm-*Hgf*-No-XHs and Mm-*Hgf*-C3, in marginal cells and to a lesser extent in the spiral ligament and Reissner’s membrane (Fig. 7*C*).

**Figure 7.**
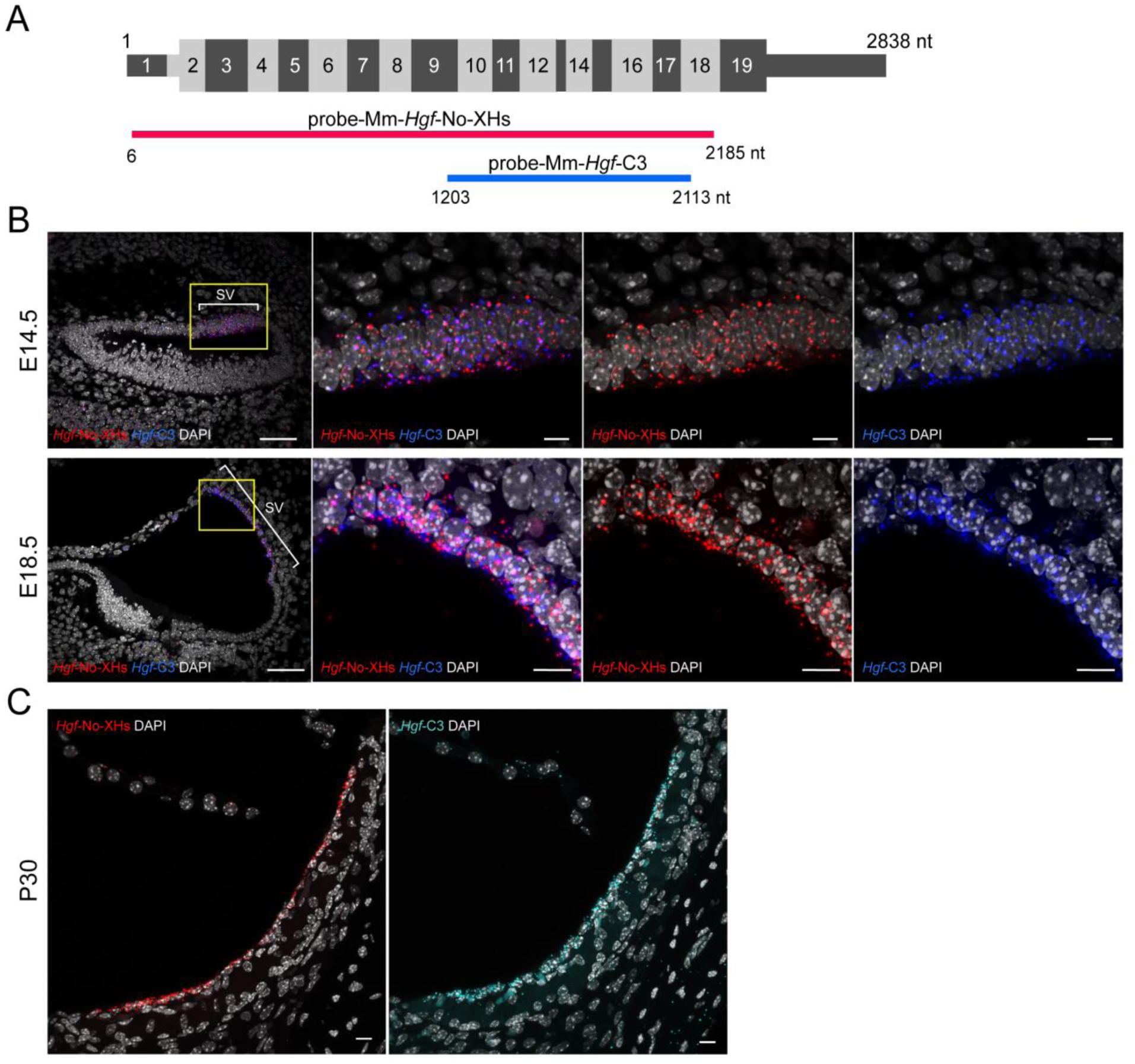
*Hgf* mRNA expression in the developing and adult stria vascularis. ***A***, Schematic showing targeted regions for smFISH RNAscope probes for *Hgf* mRNA. *Mm*-*Hgf*-No-XHs (red) corresponds to the full-length sequence of *Hgf* mRNA while *Mm-Hgf*-*C3* (blue) targets the sequence towards the C-terminus of *Hgf* mRNA. In effect, these probes enable dual methods of detecting the full-length form of *Hgf*. ***B***, Expression of *Hgf* mRNA in the perinatal stria vascularis (SV) at E14.5 and E18.5. Representative images of E14.5 and E18.5 stria vascularis co-labeled with smFISH probes for Mm-*Hgf*-No-XHs (red) and Mm-*Hgf*-C3 (blue). The full-length form of *Hgf* is detected by both probes at E14.5 and E18.5. Scalebars are 50 µm (first row) and 10 µm (other rows). ***C***. Expression of *Hgf* mRNA in the adult stria vascularis (P30). Representative images of P30 stria vascularis co-labeled with smFISH probes for Mm-*Hgf*-No-XHs (red) and Mm-*Hgf*-C3 (turquoise) demonstrate expression of *Hgf* RNA in the adult stria vascularis. The full length form of *Hgf* is detected by both probes at P30. Scalebars are 10 µm. Artifactual labeling of capillaries noted.

## Discussion

The ionic composition of cochlear endolymph within the inner ear has a notably high potassium concentration of 150 mM, which is necessary for hair cell mechano-electrical transduction of sound. In this study we demonstrate a mechanistic link between dysfunctional ion homeostasis in the inner ear and a 10bp deletion (del10) of conserved intronic sequence of *Hgf*, which accounts for the neurosensory deafness of *Hgf*^del10Neo/del10Neo^ and *Hgf* ^del10/del10^ mutant mice and, by analogy, is the likely reason for human DFNB39 deafness. Homozygosity for the del10 mutation results in a failure of neural crest-derived melanocytes to incorporate into the SV during development, leading to a reduced intermediate cell layer and consequently, compromised endocochlear potential (EP), deafness and subsequent hair cell loss. Thus, DFNB39 qualifies as a neurocristopathy that is surprisingly nonsyndromic (Bolande, 1974; Vega-Lopez et al., 2018; Ritter and Martin, 2019). Mouse models described here provide an opportunity to study the role of HGF-MET signaling in neural crest cell incorporation into the SV.

SV intermediate cells express the KCNJ10 inward-rectifying potassium channel, which is critical for EP generation (Tachibana, 1999; Marcus et al., 2002; Wangemann et al., 2004). The EP reduction in both *Hgf* ^del10Neo^ and *Hgf* ^del10^ homozygous KI mice suggests that the initial tissue affected is the SV. Cell type-specific protein expression for markers of marginal (SLC12A2), intermediate (KCNJ10), and basal (CLDN11) cells reveals a reduction in KCNJ10 protein expression in both *Hgf* ^del10Neo^ and *Hgf* ^del10^ homozygous KI mice. This is consistent with the observed reduction in future intermediate cells derived from *Dct*-expressing neural crest cells seen in the *Hgf*-CKO mouse line (Fig. 5; (Shibata et al., 2016)), and with the depletion of intermediate cell specific transcripts in homozygous mutant cochleae (Fig. 6*C*). These results link the hearing loss phenotype to an intermediate cell reduction, supporting the contention that neural crest-derived intermediate cells play a significant role in EP generation by the SV.

While SLC12A2 and CLDN11 protein levels remain unchanged between homozygous KI and wild type *Hgf* ^del10/del10^ littermates, SLC12A2 demonstrates a slight, significant increase in expression in the *Hgf* ^del10Neo^ line. Because marginal cells utilize SLC12A2 to transport potassium from the intrastrial space to maintain the low intrastrial potassium essential for EP generation (Takeuchi et al., 2000), the severity of the hearing loss in *Hgf* ^del10Neo^ mice may result in the induction of a compensatory mechanism heralded by an increase in SLC12A2 protein expression to account for the diminished intermediate cell population’s ability to generate the high potassium concentration necessary for the EP. However, a change in expression of certain proteins like SLC12A2 may upset the fine balance necessary for normal function and result in dysfunction instead of compensation. For these reasons, the effect of the Neo cassette may be a tool to explore the response of marginal cells to reduction of SV intermediate cells.

Despite ABR threshold elevations in both the homozygous *Hgf* ^del10Neo/del10Neo^ and *Hgf* ^del10/del10^ KI mice, there is an intact organ of Corti at P60, with only a dysfunctional SV due to the reduced number of neural crest-derived intermediate cells. At P90, homozygotes have outer hair cell loss and strial and spiral ligament atrophy. A reduced EP with a normally developed organ of Corti in younger KI homozygotes supports the contention that a strial deficit precedes pathogenesis of the organ of Corti, likely due to dysfunctional ion homeostasis in the cochlear endolymph (Liu et al., 2016; Huebner et al., 2019). The timeline and relationship between EP reduction and hair cell loss has not been definitively established in this study or by others. The reduction but not absence of KCNJ10 combined with the thinning of the SV seen in homozygous *Hgf* ^del10Neo^ and *Hgf*^del10^ mice suggest the possibility of two different populations of KCNJ10-expressing cells within the SV. Light and dark cells have been reported as two distinct intermediate cell types in the SV (Cable et al., 1992). Light intermediate cells are dendritic with electron-lucent cytoplasm containing numerous cell organelles indicative of synthetic activity while dark intermediate cells possess numerous melanin granules. Another possibility is that the remaining KCNJ10-expressing cells in the homozygous KI mouse from both *Hgf*^del10Neo^ and *Hgf* ^del10^ lines may be perivascular-resident macrophage-like melanocytes (PVM/Ms) as described (Zhang et al., 2012). A third non-mutually exclusive possibility is that *Hgf* ^del10^ incompletely damages the process of neural crest cells integration into the SV.

HGF expression during inner ear development must be fine-tuned, with a deficit or an excess resulting in hearing loss (Takayama et al., 1996; Schultz et al., 2009). When there is a deficit of HGF, neural crest cells fail to migrate sufficiently into the strial intermediate cell layer. The molecular mechanism by which excess HGF causes deafness is not understood (Schultz et al., 2009). An understanding of the role of HGF in cochlear development and homeostasis will require a comprehensive survey of the spatial and temporal expression of the splice isoforms of *Hgf*. The data in this paper contribute to such a study. We show by *in situ* hybridization that *Hgf* is expressed at E14.5 and E18.5 in the SV as well as during adulthood at P30 (Fig. 7). With qRT-PCR, we show the temporal expression of alternative transcripts HGF, HGF/NK1 and HGF/NK0.5 (AK142159.1) in the developing inner ear (Fig. 1*G*). The latter is of particular interest since its 3’UTR encompasses the 10bp deletion. The HGF/NK0.5 isoform is conserved in human and terminates with only 35 of 80 residues of kringle domain K1 (Fig 1*A*). The function of HGF/NK0.5 is unknown and will be a challenge to study *in vivo*. Because an arginine residue at the C-terminus of HGF/NK0.5 is the only unique sequence not found in full-length HGF protein, obtaining a specific HGF/NK0.5 antibody will be problematic (extended Fig. 1-1). The HGF/NK1 isoform includes the entire first kringle domain and can interact with the MET receptor, as can HGF/NK2, a competitive antagonist of HGF mitogenicity (Cioce et al., 1996). This isoform diversity is further complicated by the existence of an alternate splice acceptor site in exon 6, which either removes or retains 5 amino acids from the first kringle domain. Co-expression of the resulting shorter protein, HGF_723_, along with the canonical long isoform, HGF_728_, has been shown to potentiate HGF’s angiogenic effects (Pyun et al., 2010; Hahn et al., 2011). We demonstrate that homozygosity for the 10bp deletion in the adult results in reduced usage of the exon 6a splice acceptor, leaving exon 6b levels unaltered. The reduction of 6a acceptor usage was seen only in the cochlea, and not in kidney or lung (Fig. 6B). A future study will concentrate on the diversity and functional roles of each HGF isoform using mouse models that permit tissue-specific, isoform-specific and quantitative regulation of *Hgf* expression, including the NK0.5 isoform, as well as the 6a and 6b splice acceptor sites during the developmental period when neural crest cells migrate to become SV intermediate cells.

Although we have focused on the developmental component of the mutant phenotype, the persistent expression of MET in adult SV intermediate cells suggests an ongoing requirement for HGF/MET signaling after SV development is complete (Shibata et al., 2016). In the adult, the ototoxic effects of aminoglycoside antibiotics can be ameliorated by exogenous HGF or an HGF mimetic, preventing hair cell loss (Kikkawa et al., 2009; Uribe et al., 2015). Specifically, exogenous HGF appears to be protective against neomycin in cochlear explants (Kikkawa et al., 2009). Dihexa, a HGF mimetic, crosses the blood-brain barrier and protects against acute aminoglycoside ototoxicity by upregulating MET in hair cells in zebrafish neuromasts (Uribe et al., 2015). These studies suggest that when MET expression is altered, increasing HGF expression may prevent hair cell death.

In summary, we describe a mouse model of human deafness DFNB39 and demonstrate a functional link between a 10bp deletion in a highly conserved intronic sequence of *Hgf* and hereditary hearing loss. The del10 mutation diminishes *Hgf* expression specifically in the cochlea, leading to failure of neural crest derived melanocytes to infiltrate the SV during development. Consequently, the intermediate cell layer is reduced and compromised, leading to reduced endocochlear potentials, hearing loss and ultimately, hair cell loss. The *Hgf* ^del10Neo^ and *Hgf* ^del10^ lines will be invaluable resources in further studies of HGF expression and function in the auditory system.

